# RhCMV Expands CCR5 Memory T Cells and promotes SIV reservoir genesis in the Gut Mucosa

**DOI:** 10.1101/2025.01.06.631163

**Authors:** Chrysostomos Perdios, Naveen Suresh Babu, Celeste D. Coleman, Anna T. Brown, Matilda J. Mostrom, Carolina Allers, Lara Doyle-Meyers, Christine M. Fennessey, Brandon F. Keele, Amitinder Kaur, Michael L. Freeman, Joseph C. Mudd

## Abstract

Cytomegalovirus (CMV) is a prevalent β-herpesvirus that persists asymptomatically in immunocompetent hosts. In people with HIV-1 (PWH), CMV is associated with persistence of the HIV-1 reservoir and particular inflammatory related co-morbidities. The true causative role of CMV in HIV-associated pathologies remains unclear given that nearly all PWH are coinfected with CMV. In this study, we examined acute phase SIV dynamics in cohorts of rhesus macaques that were seropositive or -negative for rhesus CMV (RhCMV). We observed expansion of CCR5+ target CD4+ T cells in gut and lymph nodes (LN) that existed naturally in RhCMV-seropositive animals, the majority of which did not react to RhCMV lysate. These cells expressed high levels of the chemokine receptor CXCR3 and a ligand for this receptor, CXCL9, was systemically elevated in RhCMV-seropositive animals. RhCMV+ RMs also exhibited higher peak SIV viremia. CCR5 target memory CD4 T cells in the gut of RhCMV+ RMs were maintained during acute SIV and this was associated with greater seeding of SIV DNA in the intestine. Overall, our data suggests the ability of RhCMV to regulate chemotactic axes that direct lymphocyte trafficking and promote seeding of SIV in a diverse, polyclonal pool of memory CD4+ T cells.

## Introduction

Cytomegalovirus (CMV; HHV-5), a β-herpesvirus, commonly infects humans from a young age. The global anti-CMV IgG seroprevalence is highly prevalent with a worldwide average of 83%^1^. Only within immunocompromised settings does CMV manifest overt disease^2^. Nonetheless, maintaining CMV in an asymptomatic state presents a significant burden to the immune system. CMV-specific CD4+ and CD8+ T cells dominate the immune landscape in blood, and approximately 10% of circulating T cells can be reactive to CMV peptides in seropositive healthy adults^3^. These cells are characteristically oligoclonal and exhibit a differentiated effector memory or terminal effector phenotype^4^. As individuals age with CMV, virus-specific cells expand and comprise an increasing fraction of T cells in blood, particularly within the CD8+ T cell compartment^4^. This phenomenon, known as CD8+ memory T cell inflation, constitutes a classical aging-associated immune profile and has been linked with several aging associated morbidities including cardiovascular disease, frailty, and neurocognitive impairment^5–7^.

Co-infections are extremely common in people with HIV-1 (PWH)^8^, with CMV coinfections among the most prevalent with more than 90% of PWH being CMV-seropositive^9^. Compared to their numbers in individuals without HIV-1, CMV-specific CD4+ and CD8+ T cells are further elevated in the circulation of PWH, on par to levels observed in elderly populations, and remain elevated even with virus suppression by long-term antiretroviral therapy (ART)^10,4,11^. In PWH, the clonal expansion of CMV-specific T cells is believed to contribute to cardiovascular complications and support the persistence of the HIV-1 reservoir^12–16^. Asymptomatic CMV infection may also shape the immune system more broadly. For example, systems-based approaches in healthy CMV discordant twins without HIV-1 revealed that out of 204 innate and adaptive immune measurements, 119 (58%) were influenced by CMV serotype^17^. Many of these point to a profound influence of CMV on the cytokine and chemokine milieu, including plasma interferon-γ (IFNγ), IL-6, and IL-10 as antigen-independent immune mediators that were elevated in CMV-seropositive donors^18–20^. In PWH, co-infection with CMV is associated with substantial dysregulation of the plasma proteome^21^, including the increased production of several inflammatory cytokines and chemokines such as CXCL10 (IP-10) and soluble (s)TNF-RII^11^. Furthermore, blockade of asymptomatic CMV replication with the antiviral drug letermovir in ART-suppressed PWH was recently shown to induce broad declines in a number of inflammatory and cardiometabolic plasma proteins^22^. While these data suggest influence of CMV by mechanisms that go beyond the clonal expansion of CMV-specific cells, how this unique immunological environment can shape HIV-1 disease outcomes is less clear.

The high seroprevalence of CMV in PWH is a major obstacle to determining causal relationships. Previous studies have stratified CMV-seropositive cohorts of PWH by the presence or absence of CMV shedding in genital secretions with the expectation that genital shedding is indicative of antigenic burden^23,24^. Subclinical CMV shedding is largely intermittent however and may not fully capture the CMV-driven immunologic imprint. In this study, we assessed the impact of rhesus CMV (RhCMV) co-infection on the acute phase response to simian immunodeficiency virus (SIV) infection in a nonhuman primate model using rhesus macaques that were seronegative (RhCMV-) or seropositive (RhCMV+) for RhCMV. We observed expansion of CCR5+ target CD4+ T cells in gut and lymphoid tissues that existed naturally in RhCMV+ animals prior to SIV infection, the majority of which were not RhCMV-specific. These cells expressed high levels of the chemokine receptor CXCR3 and a ligand for this receptor, CXCL9, was systemically elevated in RhCMV+ animals. Following infection with SIV, CXCR3-expressing memory CD4 T cells were enriched in gut mucosal tissues of RhCMV+ animals, which was associated with maintenance of CCR5+ target cells in the gut and greater seeding of SIV DNA in the small and large intestine. Overall, our data suggest the ability of RhCMV to regulate chemotactic axes that direct lymphocyte trafficking and promote seeding of SIV in a diverse, polyclonal pool of memory CD4+ T cells.

## Results

### RhCMV+ animals have naturally higher percentages of CCR5+ Th1-like memory CD4 T cells in blood and tissues

In order to characterize phenotypic differences in lymphocyte populations associated with differential RhCMV infection, tissue samples from blood, peripheral lymph nodes (LN), bone marrow (BM), duodenum and colon were biopsied in 8 RhCMV- and 12 RhCMV+ healthy adult male and female rhesus macaques (Table 1). RhCMV+ animals in this study were derived from an outdoor colony in which CMV seroprevalence is endemic, acquired across mucosal surfaces within the first year of life by horizontal transfer of the virus through bodily fluids^25,26^. RhCMV-animals were derived from a specialized colony that were nursery reared or born to RhCMV-mothers and remained seronegative for RhCMV and other persistent viruses throughout life (Table 1). Seroprevalence of rhesus lymphocryptovirus (RhLCV; Macacine gammaherpesvirus 4), a persistent γ-herpesvirus that is highly homologous to human Epstein-Barr virus (EBV)^27^, was variable among RhCMV+ and RhCMV-groups (Table 1).

**Table 1:** Pre-SIV study group animal demographics.

| Characteristics | All | CMV- | CMV+ | p-value |
| --- | --- | --- | --- | --- |
| n | 21 | 9 | 12 |  |
| Sex(M/F) | 8/13 | 3/6 | 5/7 | 1 |
| Age (mean; years) | 12.50 | 12.42 | 12.56 | 0.806 |
| Age (min-max; years) | 5.77-18.06 | 6.49-18.06 | 5.77-17.75 |  |
| Weight (mean; kg) | 11.14 | 10.47 | 11.64 | 0.804 |
| Weight (min-max; kg) | 6.30-18.20 | 6.75-14.80 | 6.30-18.20 |  |
| LCV+ (%) | 81% | 66% | 92% | 0.378 |

We first determined whether latent asymptomatic RhCMV infection was associated with effector memory CD4+ and CD8+ T cell expansion in blood (gating strategy in Supp. Fig. 1), which is a hallmark immunological imprint of persistent asymptomatic CMV infection in humans^10,4^. RhCMV+ animals exhibited comparable numbers of naïve (T_N_), central (T_CM_), and transitional memory (T_TM_) T cells in blood, however CD4+ and CD8+ effector memory T cells (T_EM_) were notably expanded in RhCMV+ animals (Supp. Fig. 2A, B). These differences were not observed when animals were stratified by RhLCV-serostatus (Supp. Fig. 2C, D). We next assessed whether RhCMV was associated with particular CD4+ T cell activation states known to influence HIV-1/SIV permissiveness and productive infection. We defined expression of 3 phenotypic markers on total CD95+ memory CD4+ T cells, including the HIV-1/SIV co-receptor CCR5, the activation marker HLA-DR, and cell cycling marker Ki-67, and performed hierarchal clustering of these markers across tissues. We detected no discernable pattern of HLA-DR and Ki-67 expression that distinguished these markers by RhCMV serostatus, whereas CCR5 expression on memory CD4+ T cells clustered distinctly between RhCMV+ and RhCMV-animals, with only 2 of the total 21 animals that did not stratify by RhCMV serotype (Fig. 1A). Expression of CCR5 memory CD4+ T cells was significantly higher on memory CD4+ T cells within the blood, colon, and especially within the LN of RhCMV+ animals compared to RhCMV-animals (Fig. 1B), but not in RhLCV+ animals compared to RhLCV-animals (Supp Fig. 3A). These differences were also noted on CD8+ memory T cells within the LN and duodenum of the RhCMV+ group (Supp. Fig. 3B), but not when stratified by RhLCV serostatus (Supp. Fig. 3C). Plasma levels of two CCR5 ligands, CCL3 and CCL4, which directly bind and internalize the CCR5 receptor, were similar among RhCMV- and RhCMV+ groups (Supp. Fig. 3D, E), suggesting that RhCMV-associated increases in CCR5 surface density were not due to altered ligand-receptor occupancy. We also assessed CCR5 surface expression in CD4+ memory phenotypes in a human cohort of CMV-seropositive and -seronegative adults without HIV-1. Surface expression of CCR5 was upregulated on terminal effector memory CD4+ T cells (T_EMRA_) in blood (Supp. Fig. 3F) of these healthy human CMV-seropositive donors, suggesting similar immunologic impacts between the rhesus and human CMVs.

**Figure 1:**
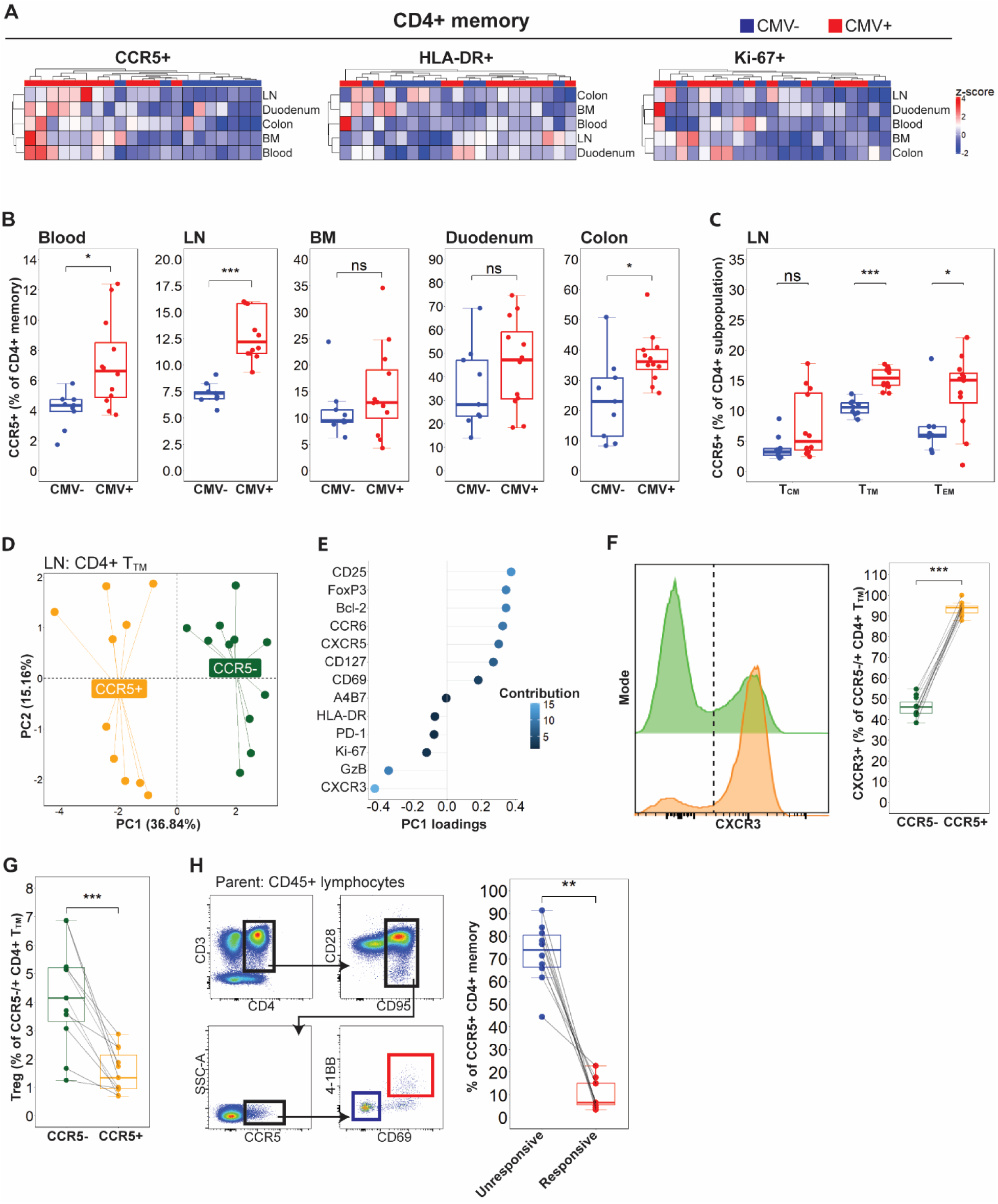
**A** Heatmaps with unsupervised clustering by CMV-serostatus on %CCR5/HLA-DR/Ki-67 CD4+ memory T cells across sampled tissues (CMV+ n = 12; CMV-n = 9). **B** %CCR5+ CD4+ memory boxplots in each sampled tissue (CMV+ n = 12; CMV-n = 9). **C** %CCR5+ across CD4+ T_CM_, T_TM_ and T_EM_ subpopulations (CMV+ n = 12; CMV-n = 9). **D** PCA plot of CCR5+ and CCR5-CMV+ CD4+ T_TM_ cells in the CMV+ LN (n = 11). **E** PC1 loadings with color coded contribution % for the PCA plot (n = 11). **F** Histogram and %CXCR3+ between CCR5+/-CD4+ T_TM_ in CMV+ LN (n = 11). **G** %Treg between CCR5+/-CD4+ T_TM_ in CMV+ LN (n = 11). **H** Gating strategy and %CCR5+ CD4+ memory cells CMV-specificity based on 4-1BB and CD69 expression (n = 10). Error bars represent 1.5 times the interquartile range. For B-C statistical comparisons performed using two-sided Mann-Whitney U test. For F-H statistical comparisons performed using two-sided Wilcoxon matched pairs signed-rank test. Key: LN, lymph node; BM, bone marrow; T_CM_, central memory; T_TM_, transitional memory; T_EM_, effector memory. ns p>0.05; * p<0.05; ** p<0.01; *** p<0.001.

Because the total CD95+ memory CD4 T cell pool in RMs contains subpopulations of varying maturation levels, we measured CCR5 expression on distinct maturation subsets within the CD4+ memory pool comprising of T_CM_, T_TM_, and T_EM_ cells. In the RhCMV+ LN where CCR5 expression was most starkly enriched, expression of CCR5 was found to be significantly higher in T_TM_ and to a lesser degree, T_EM_ CD4+ T cells, whereas this contrast was not significant in the CD4+ T_CM_ subpopulation (Fig. 1C). CCR5+ CD4+ T_TM_ cells in the LN of RhCMV+ animals were phenotypically distinct from their CCR5-counterparts and demonstrated complete separation by principal component analysis (PCA) when characterized by an array of phenotypic markers related to lineage, survival, migration, activation, and exhaustion (Fig. 1D, E). The phenotypic marker that most strongly segregated CCR5+ from CCR5-CD4+ T_TM_ cells was CXCR3, a marker that defines IFNγ producing Type-1 helper (Th1) CD4+ T cells^28,29^ (Fig. 1E), Indeed, CCR5+ CD4+ T_TM_ cells were near-uniformly positive for CXCR3 surface expression (Fig. 1F). Conversely, expression of the markers CD25 and FoxP3 were the most distinguishing features in CCR5-CD4+ T_TM_ cells, signifying enrichment of a T regulatory lineage-defining phenotype (Fig. 1G).

Because CMV-specific CD4+ T cells exhibit canonical Th1 functionalities^30,31^ , we asked whether the increase of CCR5+ target CD4+ T cells in RhCMV+ animals could be explained by an expansion of RhCMV-specific CD4+ T cells that expressed CCR5. We stimulated peripheral blood mononuclear cells (PBMCs) overnight from RhCMV+ animals with a lysate derived from RhCMV-infected fibroblasts and confirmed the specificity of CD4+ T cells reacting to RhCMV peptides by activation-induced markers 4-1BB and CD69 (Supp. Fig. 4). In RhCMV+ animals, the majority of CCR5+ memory CD4 T cells did not respond to RhCMV peptides (Fig. 1H). Taken together, the data suggest that RhCMV is associated with whole-body expansion of memory CD4+ T cells that co-express CCR5 and CXCR3, the majority of which are not RhCMV-specific.

### RhCMV is associated with higher concentrations of particular CXCR3 chemokines in circulation

To determine differences in key cytokines, and other inflammatory proteins between RhCMV+ and RhCMV-animals at baseline, we performed high-throughput proteomic profiling of over 50 analytes in plasma by proximity extension assay. Four analytes were under expressed in RhCMV+ animals relative to their levels in RhCMV-animals, including a modulator of neutrophil chemotaxis CXCL1^32^, soluble CX3CL1 (fractalkine), and soluble CD40 (Fig. 2A). CXCL9, a chemokine that binds the CXCR3 chemokine receptor^33^, was the most significantly overexpressed protein in plasma of RhCMV+ animals (Fig. 2A, B). CXCL9 was not found to be associated with RhLCV status in the smaller cohort (Supp. Fig. 5A). Levels of CXCL9 in plasma were associated with greater enrichment of CXCR3-expressing CCR5+ memory CD4+ T cells in the LN (Fig. 2C).

**Figure 2:**
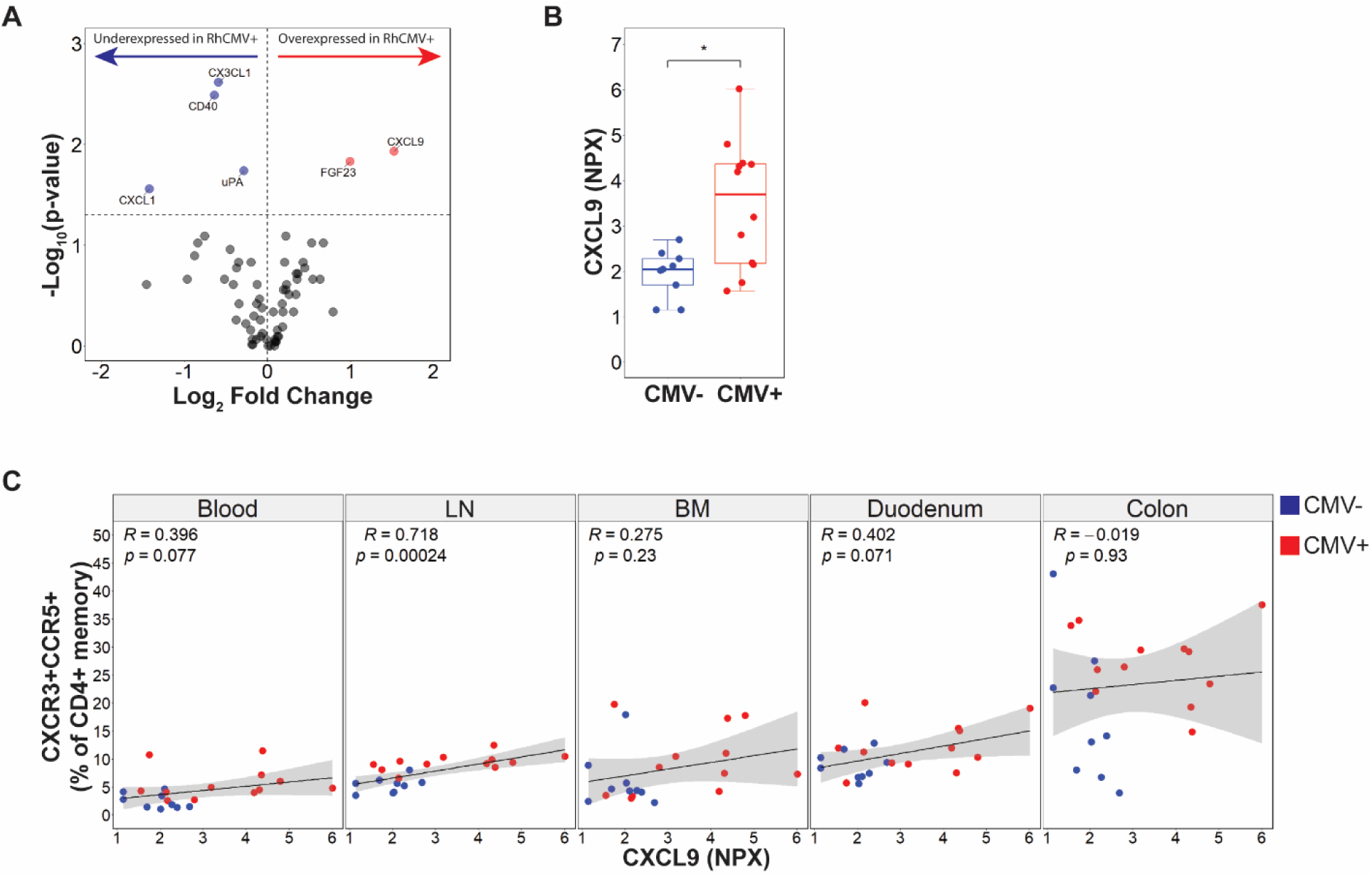
**A** Volcano plot representing the mean log_2_ fold change in plasma chemokines found in CMV+ animals (CMV+ n = 12; CMV-n = 9). Red overexpressed; blue underexpressed, gray nonsignificant. **B** CXCL9 plasma expression between CMV-serostatus. Error bars represent 1.5 times the interquartile range. Statistical comparison performed using two-sided Mann-Whitney U test. **C** Two-sided Spearman’s correlation between %CXCR3+CCR5+ CD4+ memory against CXCL9 expression (CMV+ n = 12; CMV-n = 9). Shaded area represents 95% confidence interval. Key: LN, lymph node; BM, bone marrow; NPX, normalized protein expression; * p<0.05.

**Figure 3:**
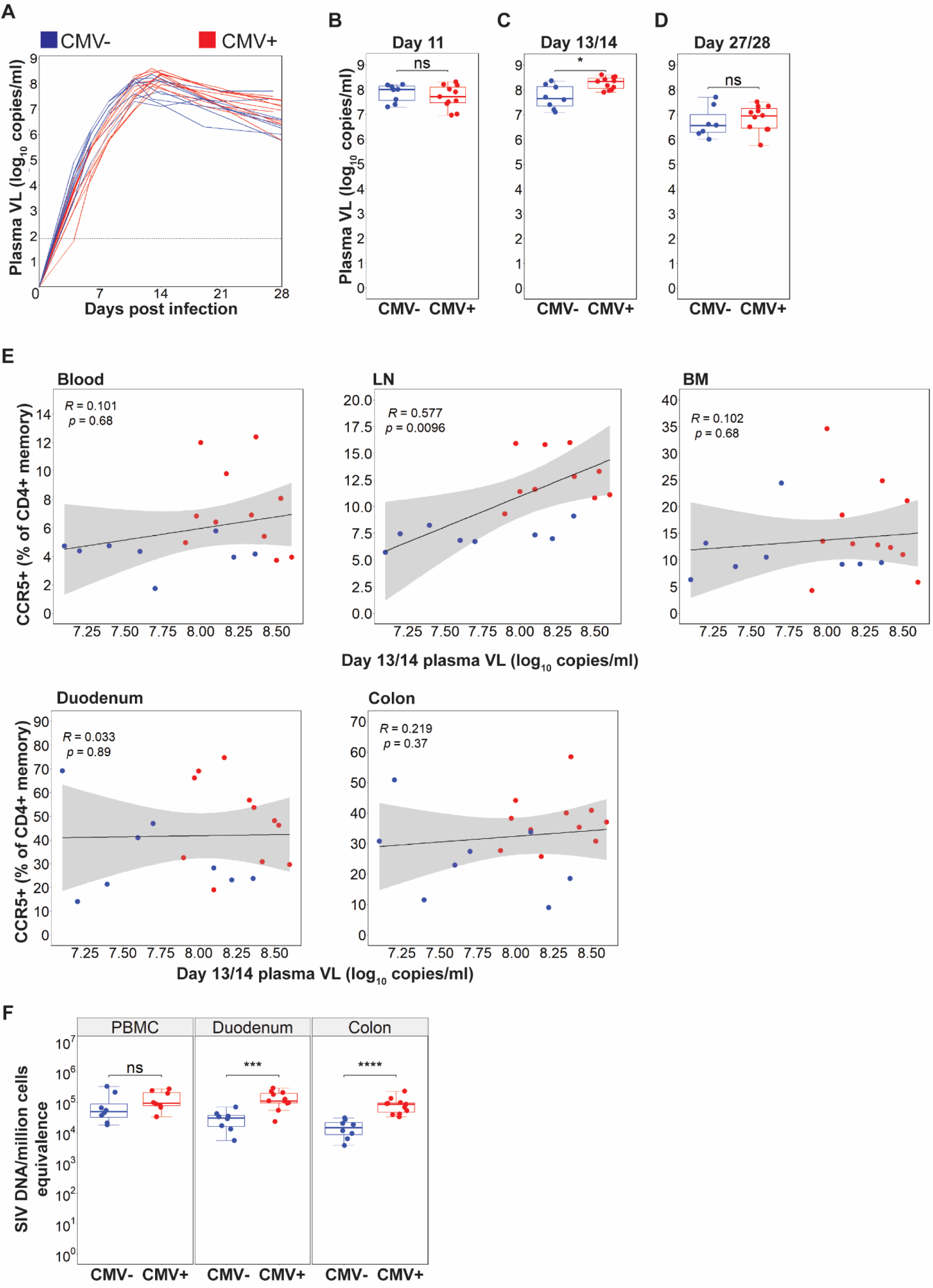
**A** Line graph of plasma SIV viral loads (VL) over days post infection. **B-D** Plasma VL on day 11, day 13/14 and day 27/28 by CMV-serostatus. **C** Two-sided Spearman’s correlation between %CCR5+ CD4+ memory against plasma VL (CMV+ n = 11; CMV-n = 8). Shaded area represents 95% confidence interval. Key: LN, lymph node; BM, bone marrow. **F** Cell-associated SIV DNA metrics by CMV-serostatus. For all graphs CMV+ n = 11; CMV-n = 8. For B-D, F error bars represent 1.5 times the interquartile range. Statistical comparison performed using two-sided Mann-Whitney U test. Key: LN, lymph node; BM, bone marrow; PBMC, peripheral blood mononuclear cells; ns p>0.05; * p<0.05; *** p<0.001; **** p<0.0001.

We further probed the association of CXCL9 with RhCMV-serostatus by examining the concentrations of this chemokine in plasma within a separate, larger cohort of RhCMV-seropositive RMs that exhibited varying frequencies of CD8+ T_EM_ expansion in blood, the hallmark immune imprint of CMV. The chemokines CXCL10 and CXL11 were also examined, which exist within a coordinated IFNγ responsive network (Supp. Fig. 5B). CD8+ T_EM_ frequencies in RhCMV+ RMs were stratified by quartile (Q1: lowest, Q4: highest) and the concentrations of CXCL9, CXCL10, and CXCL11 in plasma were compared (Supp. Fig. 5C). Both plasma CXCL9 and CXCL10 levels were highest among RhCMV+ animals falling within the quartile of greatest CD8 T_EM_ expansion (Q4; Supp. Fig. 5D-E). Although the overall comparison of CXCL11 levels across quartiles did not reach statistical significance based on the Kruskal-Wallis test, a post-hoc analysis revealed a significant difference between the lowest and highest quartiles (Supp. Fig. 5F). Taken together, the data suggest that the CXCR3/CXCL9 axis is over-represented in natural RhCMV infection and associates with greater RhCMV-driven immunological burden.

### Enhanced SIV burden in RhCMV+ animals at day 13/14 of acute SIV infection

CXCL9 is strongly induced by IFNγ and can direct Th1 cell trafficking to inflammatory sites^34–36^. In both CD4+ and CD8+ T cells, molecules that distinguish a Th1 program are commonly associated with more effective viremic control^37–39^. On the other hand, Th1 cells are readily infected by HIV-1/SIV and can harbor large clonally expanded populations of HIV-1 proviruses during ART^40,41^. To determine the relative benefit or detriment of a Th1-skewed environment that was imparted by RhCMV on acute SIV replication dynamics, we infected study animals intravenously with barcoded SIV239M and sampled whole blood at time windows of exponential SIV growth (4, 6, 8 days post-infection (dpi)), peak viremia (11, 13/14 dpi), and post-peak SIV decline (21, 28dpi; Fig. 3A). Within time windows of peak SIV replication, there were no significant differences in viremia among RhCMV- and RhCMV+ animals at 11dpi (Fig. 3B), however at 13/14dpi plasma SIV load was higher in coinfected animals (Fig. 3C). SIV viral load at 13/14 dpi was not associated with RhLCV status (Supp. Fig. 6A). Elevated viremia in this group was transient as both groups exhibited similar viral loads after post-peak SIV decline at 28dpi (Fig. 3D). We observed a direct association of peak viremia with CCR5-expressing target memory CD4+ T cells that existed prior to SIV infection in LN, but not in blood, bone marrow, or gut mucosal tissues (Fig. 3E), suggesting target cell frequencies in LN are more predictive of peak viremia than are frequencies at other anatomical sites.

We next determined the levels of total cell-associated SIV DNA in PBMCs and gut mucosal tissues at 14dpi. There were no significant differences in the frequencies of cells harboring SIV DNA within the blood of RhCMV- and RhCMV+ animals (Fig. 3F). In duodenal and colonic tissues, however, total SIV DNA levels were dramatically higher at peak viremia in co-infected animals (Fig. 3E). The RhCMV+ colon in particular exhibited median SIV DNA levels that were roughly ∼0.8 log_10_ higher than colonic tissue of the RhCMV-group (RhCMV+: 4.94; RhCMV-: 4.18; Fig. 3F). No significant differences were observed in gut cell-associated SIV DNA based on RhLCV status (Supp. Fig. 6B). Taken together, the data suggest that the impact of RhCMV/SIV coinfection on acute phase viral burden is more prominent at tissue sites than in blood, with significantly higher seeding of viral DNA occurring within gut mucosal tissues.

### CCR5-expressing memory CD4 T cells are maintained in the gut of co-infected animals during acute SIV

To examine how RhCMV influenced immune phenotypic profiles during acute SIV, we assessed surface expression of the same markers characterized pre-infection (Fig. 1) on memory CD4+ and CD8+ T cells at day 13/14 post SIV infection. Both CD4+ and CD8+ T cell counts in blood were similar among RhCMV+ and RhCMV-animals at peak viremia (13/14dpi; Fig. 4A, B). Although there were no significant differences across tissues in the fold change (FC) of CD4+ T cell frequencies at peak viremia relative to pre-infection levels, CD4+ T cell frequencies in both groups exhibited the most precipitous reduction within the colon and duodenum (Fig. 4C), in line with early target cell depletion at these sites^42,43^. Surface expression of HLA-DR was higher on blood CD4+ memory T cells of co-infected animals, as well as in the LN and BM, but not within the colon and duodenum (Fig. 4D). There were no significant differences in the percentages of Ki-67+ CD4+ memory cells in any of the tissues examined (Fig. 4E). Within the memory CD8+ T cell compartment, HLA-DR expression in co-infected animals was significantly higher in the blood, BM, and colon but not within LN and duodenal tissues (Fig. 4F). Despite an overall hyperactivated state during coinfection, frequencies of cycling CD8 T cells in the duodenum during acute SIV were notably lower in RhCMV+ animals (Fig. 4G). Lower frequencies of cycling memory CD8+ T cells in the duodenum were markedly associated with higher levels of total SIV DNA at this site (Fig. 4H), suggesting a blunted CD8+ T cell response in the duodenum during acute SIV in coinfected animals.

**Figure 4:**
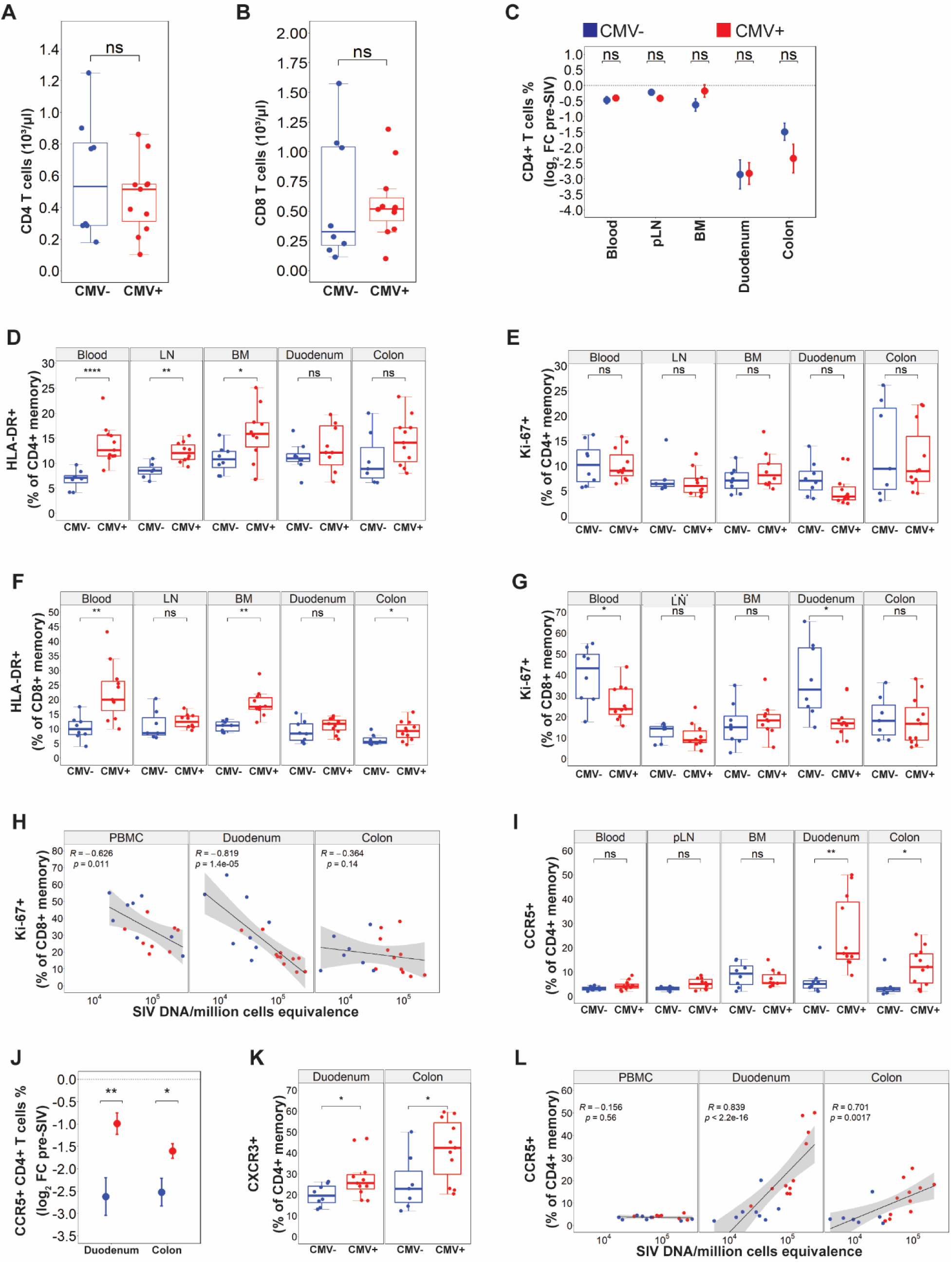
**A** CD4+ T cell numbers in blood by CMV-serostatus. **B** CD8+ T cell numbers in blood by CMV-serostatus. **C** %CD4+ T cells mean ± standard error across sampled tissues by CMV-serostatus. **D** %HLA-DR+ CD4+ memory cells across sampled tissue by CMV-serostatus. **E** %Ki-67+ CD4+ memory cells across sampled tissue by CMV-serostatus. **F** %HLA-DR+ CD8+ memory cells across sampled tissue by CMV-serostatus. **G** %Ki-67+ CD8+ memory cells across sampled tissue by CMV-serostatus. **H** Two-sided Spearman’s correlation between %Ki-67+ CD8+ memory against cell-associated SIV DNA. **I** %CCR5+ CD4+ memory cells across sampled tissue by CMV-serostatus. **J** %CCR5+ CD4+ T cells mean ± standard error across duodenum and colon by CMV-serostatus. **K** %CXCR3+ CD4+ memory across duodenum and colon by CMV-serostatus. **L** Two-sided Spearman’s correlation between %CCR5+ CD4+ memory cells against cell-associated SIN DNA. For all figures CMV+ n = 11, CMV-n = 8. For A, D-G, I, K-L error bars represent 1.5 times the interquartile range. Statistical comparison performed using two-sided Mann-Whitney U test. For H, I, L shaded area represents 95% confidence interval. Key: LN, lymph node; BM, bone marrow; PBMC, peripheral blood mononuclear cells; ns p>0.05; * p<0.05; ** p<0.01; **** p<0.0001.

We also probed how coinfection influenced phenotypic profiles of circulating monocytes at 13/14 days post-SIV. The type I IFN-stimulated surface marker CD169 was found to be elevated on monocytes of RhCMV+ animals (Supp. Fig. 7A), a finding concurrent with higher concentrations of IFNα at peak periods of plasma viremia (Supp. Fig 7B). Frequencies of CD169+ monocytes positively correlated with day 13/14 viremia (Supp. Fig 7C), together suggesting a more robust type I IFN response during acute SIV in coinfected animals.

Lastly, we assessed CCR5+ target cells across tissues during acute SIV and how their frequencies were influenced by RhCMV coinfection. CCR5+ memory CD4 T cells within blood and LNs were equally reduced among RhCMV- and RhCMV+ animals at 13/14 days post-SIV infection (Fig. 4I). At gut mucosal sites, however, RhCMV+ animals continued to exhibit expanded target cell frequencies in the colon during acute SIV (Fig. 4I). Frequencies of target cells were also elevated in the duodenum of coinfected animals (Fig. 4I), a feature that was not observed at this site prior to SIV (Fig. 2). Relative to target cell frequencies in the gut pre-SIV, RhCMV-animals exhibited greater depletion of target cells at gut mucosal tissues than that of RhCMV+ animals (Fig. 4J), despite the latter group overall exhibiting more local SIV burden (Fig. 3F).

We sought to determine potential mechanisms underlying the apparent discordant finding of greater SIV DNA seeding (Fig 3F), yet less target cell reduction in the RhCMV+ gut (Fig. 4J). We first considered the possibility that memory CD4+ T cells in the RhCMV+ gut exhibited superior survival capabilities that would allow them to withstand cell death during acute SIV. We examined expression of the interleukin-7 receptor CD127 and anti-apoptotic molecule Bcl-2, known to promote T cell survival^44,45^, yet found no differences in memory CD4+ T cell expression of these markers at gut mucosal sites during acute SIV by RhCMV serostatus (Supp. Fig. 8A, B). We next considered the possibility that CCR5+ target cell depletion could be offset by a greater rate of influx of memory CD4+ T cells into the RhCMV+ gut during acute SIV. RhCMV+ animals during acute SIV exhibited higher frequencies of memory CD4+ T cells in duodenal and colonic tissues that expressed the chemokine receptors CCR6 (Supp. Fig. 8C) and CXCR3 (Fig. 4K), which regulate homeostatic and inflammatory chemotaxis to gut mucosal tissues, respectively^46–48^. CXCR3 expression was not found to be significantly different in memory CD4+ T cells by CMV-serostatus in the gut pre-infection (Supp. Fig. 8D). In addition, CCR5+, but not CCR5-, CD4+ memory cells were found to harbor higher CCR6 expression in RhCMV+ RMs (Supp. Fig. 8E), despite no significant difference in plasma CCL20 expression (Supp. Fig. 8F). The enrichment of target cells in gut mucosal tissues during acute SIV correlated highly with the seeding of SIV DNA at these sites (Fig. 4L). Taken together, our data suggest that RhCMV can influence chemotactic axes that regulate lymphocyte trafficking and promote the maintenance of target cells at early sites of SIV replication in the gut.

### RhCMV+ animals during acute SIV overexpress inflammatory-related proteins associated with IFNγ

To gain insights into how latent asymptomatic RhCMV infection influenced the systemic cytokine and chemokine milieu during acute SIV, we performed similar proteomic profiling of plasmas during the window of peak viremia. No profiled soluble proteins were significantly over-expressed in RhCMV-animals (Fig. 5A). RhCMV+ animals at 13/14 dpi overexpressed 8 inflammatory-related proteins (Fig. 5A). Notable of these were CCL8, a chemokine with affinity for the CCR5 receptor^49^, and soluble PD-L1, the receptor of which (PD-1) is up-regulated on activated and functionally exhausted T cells^50,51^ (Fig. 5A). The CXCR3 chemokine CXCL9 remained elevated in plasma of coinfected animals during the window of peak SIV viremia (Fig. 5A, B), and correlated positively with both viremia and levels of viral DNA in blood and gut mucosal tissues (Fig. 5C, D). We next performed STRING analysis to infer interactions among over-expressed proteins in RhCMV animals during acute SIV. Many of these proteins were found to be associated with one another with high confidence (Fig. 5E), exhibiting high reliability of predicted/known protein-protein interactions by multiple evidence. Of note, the canonical Th1 cytokine IFNγ was predicted to occupy a central node within this network (Fig. 5F). Measuring IFNγ plasma levels by ELISA revealed that IFNγ was indeed higher in coinfected animals at 13/14 dpi (Fig. 5F). Overall, the cellular and cytokine/chemokine data are consistent with RhCMV driving an IFNγ-directed Th1 signature, which becomes more pronounced during acute SIV and is associated with greater viral burden particularly within gut mucosal tissues.

**Figure 5:**
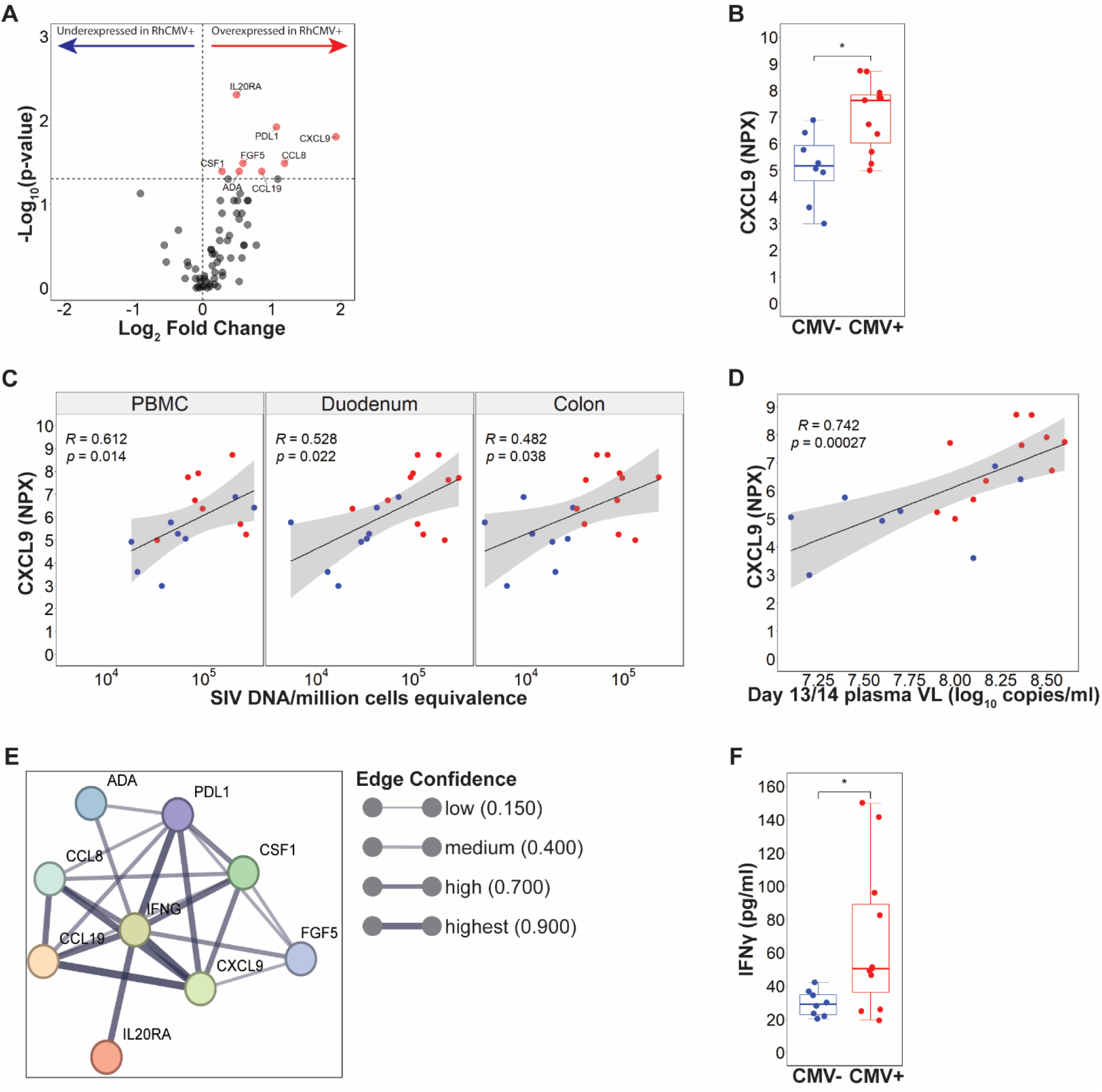
**A** Volcano plot representing the mean log_2_ fold change in plasma chemokines found in CMV+ animals. Red overexpressed; blue underexpressed, gray nonsignificant. **B** CXCL9 plasma expression between CMV-serostatus. **C** Two-sided Spearman’s correlation between plasma CXCL9 against cell-associated SIV DNA. **D** Two-sided Spearman’s correlation between plasma CXCL9 against plasma SIV viral load. **E** STRING network representing the interaction of the CMV-associated overexpressed proteins with IFNγ. **F** Plasma IFNγ by CMV-serostatus. For all figures CMV+ n = 11, CMV-n = 8. For B, F error bars represent 1.5 times the interquartile range. Statistical comparison performed using two-sided Mann-Whitney U test. For C-D shaded area represents 95% confidence interval. Key: PBMC, peripheral blood mononuclear cells; ns p>0.05; * p<0.05.

## Discussion

The establishment of HIV-1/SIV in the host and subsequent proviral burden are critically dependent on the density of CCR5-expressing target cells. In both HIV-1 and SIV, there are natural settings in which the host exhibits exceptionally low densities of target cells and these instances are associated with either spontaneous viremic control or long-term non-progression^52–55^. Here, we describe an apparent contrary setting in which natural chronic asymptomatic infection with the highly prevalent nonhuman primate homologue of human CMV leads to expansion of CCR5-expressing target cells and leads to greater seeding of SIV DNA at prominent early sites of SIV replication. This feature was not observed when stratifying our study animals by RhLCV serostatus. Given the well-documented role of CMV in influencing systemic and tissue microenvironments^56–60^, our findings suggest that RhCMV infection can alter the host immune environment in a way that enhances acute SIV replication.

We show here that target cells expanded in RhCMV+ animals exhibit distinguishing phenotypic signatures including a near uniform CXCR3+ Th1-like profile and a predominant transitional memory (T_TM_) maturation phenotype. In virally-suppressed PWH on ART, large clonal expansions of intact HIV-1 proviruses in blood can be concentrated in functionally-polarized Th1 cells^40^. Moreover, the predominant T_TM_ phenotype suggest infection of these cells could result in the seeding of SIV DNA in a maturation subtype that is long-lived, responsive to homeostatic cytokines, and known to contribute substantially to the pool of latent HIV-1 in long-term ART-suppressed persons^61^. An additional feature of target cells in RhCMV+ animals was that the majority of them did not exhibit *ex vivo* responsiveness to RhCMV-derived peptides, suggesting an apparent polyclonal repertoire extending beyond CMV-specific cells. Our assay defined RhCMV-specific CD4+ T cells by up-regulation of cognate-antigen induced surface molecules, and it is possible that some CCR5-expressing CD4+ T cells in the assay did not functionally respond to the lysate despite TCR-specificity to RhCMV epitopes. Despite this caveat, we posit that a significant fraction of target cells expanded by RhCMV are indeed polyclonal and comprise broad TCR specificities. We note that in RhCMV+ animals target cell expansion was most pronounced in lymphoid tissues. In humans, the majority of CMV-specific T cells however are excluded from lymphoid tissues and do not traffic to these sites^62–64^. At present, precisely how RhCMV expands target cells in an apparent polyclonal manner is not known, but likely related to the unique microenvironment shaped by replicating RhCMV at a persistent, low level throughout life^65,66^. Regardless of the mechanism, coinfection with RhCMV could have the potential to promote the seeding of SIV DNA in a diverse array of CD4 clonotypes beyond those that are RhCMV-specific.

We show here that expansion of CXCR3+ Th1-like target cells in RhCMV is accompanied by higher expression of the IFNγ-inducible CXCR3-binding chemokine CXCL9 in plasma. No differences in plasma CXCL9 expression in our study animals were observed when stratifying by RhLCV status. Moreover, we confirm in a separate cohort a significant trend of increasing plasma CXCL9 when RhCMV+ animals were stratified by effector CD8 T cell proportions in blood, a hallmark immunological imprint of HCMV and RhCMV^67^. In the confirmatory cohort this extended to other IFNγ-induced CXCR3-binding chemokines as well, including CXCL10 and CXCL11. The relevance of our findings to humans is supported by the observation that plasma levels of CXCL10 are elevated among ART-suppressed HCMV+ PWH compared to HCMV-PWH on ART^11^. Additionally, plasma levels of CXCL10 are elevated in lung transplant recipients with detectable HCMV replication^68^. Thus, induction of CXCR3-binding chemokines, likely existing within a larger IFNγ-regulated network, may represent a conserved mechanism by which RhCMV and HCMV influence host immunity.

RhCMV/SIV coinfection was associated with several notable acute phase virological distinctions, including transient elevation of viremia at peak SIV and starkly higher cell associated (CA)-SIV DNA frequencies in the gut. The gastrointestinal tract is a prominent site of pathology and early HIV-1/SIV replication^42,43^. Lamina propria CD4+ T cells are highly permissive to HIV-1/SIV infection and there is a near complete loss of CD4+ T cells at this site during acute infection^42,43^. In untreated HIV-1/SIV infection viral replication is anatomically dispersed, and an outstanding question is how local HIV-1/SIV replication in the gut contributes to the overall dynamics of acute phase viremia relative to other anatomic sites. In our study, it is interesting to note that at 14 days post-infection, gut mucosa CA-SIV DNA differed by near log-fold levels, yet distinctions in plasma viremia were relatively modest between RhCMV+ and RhCMV-groups and were not sustained at later timepoints of acute SIV. Our study did not employ longitudinal sampling of gut mucosal sites; however, the data suggest that at least during peak SIV, local viral replication within the gut is not a major contributor to overall levels of plasma viremia. Supporting this notion is our finding that pre-infection frequencies of target cells in the duodenum and colon did not predict levels of peak viremia. Only in lymphoid tissues were pre-infection target cell frequencies highly correlated to plasma SIV at 14 dpi. Thus, levels of peak viremia may largely be determined by the degree of viral replication in lymphoid tissues, with other anatomical sites representing a minority contribution. It is possible that this dynamic may also extend to the early emergence of viremia following analytical treatment interruptions of ART, as early recrudescing strains of SIV in plasma can be tracked specifically to CA-SIV clonotypes in lymphoid tissues^69^.

What could explain the stark enrichment of CA-SIV DNA in the gut, but not blood of RhCMV+ animals? We show here at 14 days post-infection that despite a generally higher RhCMV-associated SIV burden, frequencies of gut mucosal target cells were less depleted in RhCMV+ animals when compared to those of the RhCMV-group. Maintenance of target cells could not be explained by differing phenotypic survival signatures known to promote resistance to HIV-1/SIV-cell mediated death^70,71^, rather, both plasma CXCL9 levels and CXCR3+ memory CD4+ T cell frequencies remained elevated in the RhCMV+ gut, suggesting a degree of target cell influx that was enhanced in coinfected animals. Frequencies of CCR6-expressing memory CD4+ T cells were also elevated in the gut of coinfected animals, highlighting a potential additional mechanism of gut mucosal CD4+ T cell influx^72^. Lastly, we found that CCL8 was also highly overexpressed in RhCMV+ RMs during acute SIV, indicating a third potential migratory axis, since CCL8 can bind and induce chemotaxis of CCR5+ T cells^73^.

There are notable limitations to our study. First, we cannot infer which mechanism of target cell migration to the gut is dominant in coinfected animals, as our data suggest the contribution of multiple chemotactic routes. Second, while we highlight an apparent lack of RhLCV-mediated virological and immunological imprint in our study animals, we cannot rule out the potential influence of other persistent viruses. Unlike RhCMV-seronegative enhanced Specific Pathogen Free (eSPF) colonies, both Simian Foamy Virus (SFV) and Rhesus Rhadinovirus (RRV) are endemic in conventionally-housed groups with seroprevalence >90%^74,75^. While asymptomatic in healthy captive macaques, both viruses can mediate disease in settings of SIV-induced immunodeficiency, with SFV in particular exacerbating viremia and CD4+ T cell decline during late stages of SIV infection^76,77^. Third, it is currently unclear how these findings extend to latent CMV infection in humans. We show here in a small cohort of adults without HIV-1 that CCR5 surface expression is marginally increased on terminally differentiated effector memory (T_EMRA_) CD4+ T cells in blood of HCMV-seropositive donors. CD4+ T_EMRA_ cells exhibit the shortest half-life of all memory subsets (<3.1 months)^78,79^, and furthermore do not survive for longer than 2 days if productively infected^80,81^. The clinical significance of CD4+ T_EMRA_ cells as targets for HIV-1 is thus unclear given that these cells are labile and do not comprise the stable reservoir during long-term ART^82^. In our study animals, LN exhibited the most marked increase in CCR5 densities associated with RhCMV. Thus, evaluating target cells in lymphoid tissues if HCMV-seropositive and HCMV-negative individuals poses clinical meaningful implications for HIV-1 pathogenesis, persistence, and transmission.

In conclusion, this study provides new insights into the role of RhCMV infection in shaping the immune landscape in a way that promotes SIV replication and persistence. Our findings underscore the importance of CMV as a cofactor in HIV/SIV pathogenesis, particularly through its influence on CCR5+ CD4+ memory T cell populations and the associated inflammatory milieu. Future studies should explore the potential benefits of CMV-targeted therapies in reducing HIV/SIV reservoir seeding and persistence, which could have significant implications for the understanding of CMV/HIV-1 coinfection on reservoir persistence.

## Methods

### Animals and infection procedure

This study was designed with both groups, RhCMV- and RhCMV+, having roughly a similar number of male and female animals to avoid sex as a confounding variable. All Indian origin rhesus macaques (*Macaca mulatta*) involved in the study were bred within the Tulane National Primate Research Center (TNPRC) except three (RF73-75), who were bred and transferred from the Oregon National Primate Research Center (ONPRC). All animals were negative for the Mamu alleles commonly associated with effective SIV control, specifically Mamu-A*01, Mamu-B*08 and Mamu-B*17, except for one RhCMV-(KN94), which was positive for Mamu-B*17. Animals housed in the Specific Pathogen Free (SPF) colony naturally acquire CMV infection by the age of one year, whereas those in the enhanced Specific Pathogen Free (eSPF) colony remain CMV-free. Nine RhCMV- and 12 RhCMV+ rhesus macaques were examined at baseline of which eight RhCMV- and 11 RhCMV+ animals were infected with SIV. The animals were intravenously infected with 1 ml of SIVmac239M, containing a concentration of 5,000 IU/ml. Throughout the infection process, animals were anesthetized using telazol tiletamine hydrochloride and zolazepam hydrochloride (5 to 8 mg/kg intramuscular; Tiletamine–zolazepam, Zoetis, Kalamazoo, MI), along with buprenorphine hydrochloride (0.03 mg/kg) for pain management. From the TNPRC colony animals 35 RhCMV- and 75 RhCMV+ were randomly selected during a semi-annual health assessment (SAHA) for an extended blood draw to acquire PBMCs for CMV- specific T cells screening and plasma for Olink proteomics.

### Tissue processing

PBMCs isolation was performed using Ficoll-Paque PLUS (cytiva) gradient separation and standard procedures. Lymph nodes, duodenum and colon pinches were mechanically disrupted using gentleMACS C Tubes and gentleMACS Dissociator (Miltenyi Biotec) and filtered through a 100-μm cell strainer. Bone marrow was enriched for lymphocytes using Ficoll-Paque PLUS gradient separation. Cells were cryopreserved in 10% dimethyl sulfoxide (DMSO; EMD MiIlipore) in fetal bovine serum (FBS; Corning) and stored in liquid nitrogen.

### Complete blood count

Hematological analysis was performed at the TNPRC clinical lab. Hematological analysis was performed on whole blood, collected in EDTA tubes, using Sysmex NX-V-1000 Hematology Analyzer. The full blood count included absolute quantification, of red blood cells, neutrophils, monocytes, lymphocytes, basophils, and eosinophils.

### SIV plasma viral load quantification

Blood specimens from SIV-infected rhesus macaques were collected in EDTA anticoagulant and centrifuged at 900 x g for 15 minutes. Plasma was aspirated, aliquoted into cryovial tubes, and stored at -80°C. SIV plasma viral loads were quantified by TNPRC’s Center Pathogen Detection Core (PDQC) as previously described^83^. Briefly, SIV target cDNA and exogenous control cDNA were assayed in duplicate. QuantStudio 12k Flex (Thermo Scientific) was used with a program of 40 cycles at 95 °C for 15 s and 60 °C for 1 minute. The following primers and probe for SIVmac239 were used: Forward 5′ - AGGCTGCAGATTGGGACTTG - 3′, Reverse 5′ - TGATCCTGACGGCTCCCTAA - 3′, and Probe 5′ - FAM- ACCCACAACCAGCTCCACAACAAGGAC-IABKFQ - 3′.

### Cell-associated SIV DNA quantification

Cell-associated SIV DNA was quantified from PBMCs using Buffer RLT Plus (Qiagen; catalog 1053393) with 10% β-mercaptoethanol (Millipore Sigma; catalog M3148). Cell lysates were homogenized using QIAshredder spin columns (Qiagen, Cat. No. 79656), and the DNA was subsequently isolated using the AllPrep DNA spin column kit (Qiagen, Cat. No. 80204). Nested PCR was performed using the following primers: SIVnestF 5′ - GATTTGGATTAGCAGAAAGCCTGTTG - 3′, SIVnestR 5′ - GTTGGTCTACTTGTTTTTGGCATAGTTTC - 3′. The PCR conditions consisted of an initial 94 °C for 2 minutes, followed by 12 cycles of 94 °C for 15 seconds, 60 °C for 15 seconds, and 68 °C for 15 seconds Quantitative RT-PCR was performed on a QuantStudio 6 Flex system (Thermo Scientific) with the following cycling parameters: 50 °C for 2 minutes, followed by 40 cycles at 95 °C for 20 seconds and 60 °C for 20 seconds. The following primers were used: sGAGF 5′ - GTCTGCGTCATCTGGTGCATTC - 3′, sGAGR 5′ - CACTAGGTGTCTCTGCACTATCTGTTTTG - 3′, sGAGPr 5′ - FAM-CTTCCTCAG-ZEN-TGTGTTTCACTTTCTCTTCTGCG-3IABkFQ - 3′, CCR5F 5′ - ATGGACTATCAAGTGTCAAGTCCAACC - 3′, CCR5R 5′ - GGCGGGCTGCGATTTGTTTC - 3′ and CCR5Pr 5′ - FAM- TTTGGCAGG/ZEN/GTTCCGATGTATAATAATCGATGTC-3IABkFQ - 3′.

### Proximity extension assay (Olink)

Plasma proteins from rhesus macaques were analyzed using a proximity extension assay (Olink Proteomics). Plasma was collected and stored as described above. The Olink^®^ Target 96 inflammation panel (Olink Proteomics AB, Uppsala, Sweden) was used to measure protein levels following manufacturer’s instructions. In brief, pairs of oligonucleotide-labeled antibody probes are mixed with plasma to allow binding to their protein targets. The oligonucleotide pairs hybridize when the two compatible probes are in proximity. The reaction mixture, containing DNA polymerase, allows proximity-dependent DNA polymerization and the creation of a unique PCR target sequence, the amplified DNA sequence is quantified. Protein levels are expressed as arbitrary log_2_ transformed units, normalized protein expression (NPX).

### ELISAs

IFNγ plasma levels were measured using Rhesus Macaque IFN-gamma ELISA Kit (invitrogen; catalog: EP8RB). The essay was conducted according to manufacturer’s instructions using undiluted samples.

IFNα plasma levels were measured using Non-human IFN-alpha ELISA Kit (novusbio; catalog: NBP3-11725). The essay was conducted according to manufacturer’s instructions using 1:2 diluted samples.

### Phenotyping characterization by flow cytometry

Thawed, unstimulated rhesus macaque PBMCs, LN, bone marrow, duodenum, and colon samples were stained to identify lymphocytes and monocytes. The samples were stained in predetermined optimal concentrations of the following antibodies: BB515 anti-Siglec-1 (CD169; BD Horizon Custom; clone 7-239; catalog 624279), BB630 anti-CD28 (BD Horizon Custom; clone CD28.2; catalog 624294), BB660 anti-CCR7 (CD197; BD Horizon Custom; clone 3D12; catalog 624295), BB700 anti-CD69 (BD OptiBuild; clone FN50; catalog 747520), BB790 anti-CXCR3 (CD183; BD Horizon Customs; clone 1C6/CXCR3; catalog 624296), BV421 anti-Granzyme B (BD Horizon; clone GB11; catalog 563389), BV510 LIVE/DEAD Fixable Aqua Dead Cell Stain (Invitrogen; catalog L34957), BV605 anti-CD14 (BioLegend; clone M5E2; catalog 301834), BV650 anti-PD-1 (CD279; BioLegend; clone EH12.2H7; catalog 329950), BV711 anti-HLA-DR (BD Horizon; clone G46-6; catalog 563696), BV750 anti-CD8a (BioLegend; clone SK1; catalog 344756), BV786 anti-CD16 (BD Horizon; clone 3G8; catalog 563690), BUV395 anti-CD3 (BD Horizon; clone SP34-2; catalog 564117), BUV496 anti-CD95 (BD Horizon Custom; clone DX2; catalog 624283), BUV563 anti-Bcl-2 (BD Horizon Custom; clone Bcl-2/100; catalog 624284), BUV615 anti-CCR6 (CD196; BD OptiBuild; clone 11A9; catalog 751515), BUV661 anti-CD25 (BD OptiBuild; clone 2A3; catalog 741685), BUV737 anti-CCR5 (CD195; BD OptiBuild; clone 3A9; catalog 748873), BUV805 anti-CD4 (BD OptiBuild; clone OKT4; catalog 750976), PE anti-FOXP3 (BioLegend; clone 206D; catalog 320108), PE-eFluor610 anti-CXCR5 (CD185; Invitrogen; clone MU5UBEE; catalog 61-9185-42), PE-Cy5 anti-CD127 (Invitrogen; clone eBioRDR5; catalog 15-1278-42), PE-Cy7 anti-α4β7 (NHPRR; clone A4B7R1; catalog PR-1427); APC anti-NKG2a (CD159a, Beckman Coulter; clone Z199; catalog A60797), R718 anti-Ki-67 (BD Horizon; clone B56; catalog 566963), APC-H7 anti-CD45 (BD Pharmingen Custom; clone D058-1283; catalog 624347). Foxp3/Transcription Factor Staining Buffer Set (eBioscience; catalog 00-5523-00) was used for permeabilization for Granzyme B, Bcl-2, FOXP3, and Ki-67. The samples were stained at 37 °C for 15 min for the live/dead stain followed by 20 min for the surface antibodies and at 4 °C for 30 min for the intracellular antibodies. The stained samples were fixed in 1% paraformaldehyde (PFA) and acquired on a BD Symphony A5 cytometer. Analysis was performed using FlowJo (version 10.10.0).

### Phenotypic analysis of human PBMC

Thawed, unstimulated human PBMCs were stained to identify CCR5 expression. The samples were stained in predetermined optimal concentrations of the following antibodies: BUV395 anti-CD4 (BD Horizon; clone L200; catalog 564107), BUV737 anti-CD3 (BD Horizon; clone UCHT 1; catalog 612750), BV605 anti-CD8 (BD Horizon; clone SK1; catalog 564116), BV650 anti-CD45RO (BD Horizon; clone UCHL1; catalog 563750), BV711 anti-CCR5 (BD Horizon; clone 2D7; catalog 563395) and BV510 LIVE/DEAD Fixable Aqua Dead Cell Stain (Invitrogen; catalog L34957). The samples were stained at 37 °C for 15 min for the live/dead stain followed by 20 min for the antibodies. The stained samples were fixed in 1% PFA and acquired on a BD Fortessa cytometer. Analysis was performed using FlowJo (version 10.10.0).

### AIM assay

Thawed rhesus macaque PBMCs were stimulated with 3 µl CMV-lysate (acquired by the ONPRC virology core) at 37 °C for 16 hours. The samples were stained in predetermined optimal concentrations of the following antibodies: BB515 anti-PD-L1 (CD274; BD Horizon; clone MIH1; catalog 564554), PCP-Cy5.5 anti-CD4 (BD Pharmingen; clone L200; catalog 552838), APC anti-CD20 (BioLegend; clone 2H7; catalog 302310), APC anti-NKG2A (CD159a, Beckman Coulter; clone Z199; catalog A60797), APC-H7 anti-CD45 (BD Pharmingen Custom; clone D058-1283; catalog 624347), BV421 anti-CXCR3 (CD183; BD Horizon; clone 1C6/CXCR3; catalog 562558), BV510 LIVE/DEAD Fixable Aqua Dead Cell Stain (Invitrogen; catalog L34957), BV605 anti-CD28 (BD Horizon; clone CD28.2; catalog 562976), BV650 anti-PD-1 (CD279; BioLegend; clone EH12.2H7; catalog 329950), BV711 anti-HLA-DR (BD Horizon; clone G46-6; catalog 563696), PE anti-CD71 (BD OptiBuild; clone OX-26; catalog 744417), PE-CF594 anti-CD69 (BD Horizon; clone FN50; catalog 562645), PE-Cy5 anti-CD95 (BioLegend; clone DX2; catalog 305610), PE-Cy7 anti-4-1BB (CD137; BioLegend; clone 4B4-1; catalog 309818), BUV395 anti-CD3 (BD Horizon; clone SP34-2; catalog 564117) and BUV737 anti-CCR5 (CD195; BD OptiBuild; clone 3A9; catalog 748873). The samples were stained at 37 °C for 15 min for the live/dead stain followed by 20 min for the antibodies. The stained samples were fixed in 1% PFA and acquired on a BD Fortessa cytometer. Analysis was performed using FlowJo (version 10.10.0).

### Statistical analysis

R^84^ (v 4.2.2) was used to perform statistical analysis and graph creation. The packages ggpubr (version 0.6.0), ggplot2^85^ (version 3.5.0), ComplexHeatmap^86^ (version 2.18.0) and vegan^87^ (version 2.6.4). Protein-protein interaction analysis was performed using STRING (Search Tool for the Retrieval of Interacting Genes/Proteins; version 12.0)^88^. Wilcoxon matched pairs signed-rank test was used for non-parametric paired analysis, Mann-Whitney U test for non-parametric unpaired analysis, Spearman’s rank correlation for non-parametric correlation, and PERMANOVA for non-parametric ANOVA with permutations. Statistical significance is indicated by p < 0.05. All statistical tests are two-sided.

### Study approval

The study underwent review and approval by the institutional Animal Care and Use Committee of Tulane University. Animal care procedures adhered to the guidelines outlined in the NIH’s “Guide for the Care and Use of Laboratory Animals”. Handling procedures and containment protocols for animals were approved by the Tulane University Institutional Biosafety Committee, ensuring compliance with BSL-2 containment standards. Furthermore, the Tulane National Primate Research Center holds full accreditation from the Association for Assessment and Accreditation of Laboratory Animal Care (AAALAC), underscoring its commitment to maintaining high standards of animal welfare and research ethics. Human studies were approved by the Institutional Review Board of University Hospitals Cleveland Medical Center (IRB # 01-98-55).

## Supporting information

Supplemental metafile

## Data availability

The authors declare that data supporting the results and conclusions of this study are available within the paper and its supplementary information. Additional data that support the findings of this study are available from the corresponding authors upon reasonable request. Codes used for data analysis are found here: https://github.com/chrysperdios/. Source data are provided with this paper.

## Author Contributions

J.C.M. designed the animal study. C.P. and J.C.M. designed the laboratory study. C.P., C.D.C., N.S.B., A.T.B. and participated in tissue acquisition and processing. CMF and BFK provided SIVmac239M for study infection. C.P., C.D.C., J.C.M and N.S.B. performed experiments. C.P. and J.C.M analyzed and interpreted data. C.P. and J.C.M wrote the manuscript. M.L.F. acquired and analyzed human samples. J.C.M., C.D.C, M.L.F., N.S.B. and A.T.B. provided critical and substantive intellectual editing. C.P. and J.C.M. prepared manuscript figures. All authors approved the manuscript.

## Acknowledgements

The authors thank Brian Clagett and Xi Su for excellent technical assistance. This study was supported by the NIH-funded base grant to the TNPRC P51 OD011104 and the NIH grants R01 AI167644 and R21 OD031229. NIH S10 OD026800 was awarded to support the TNRPC Flow Cytometry Core. PDQC was funded by base grant OD011104 from the NIH. The following Research Resource Identifier (RRID):SCR supported core facilities were utilized by this study: Flow Cytometry Core (024611) and Pathogen Detection and Quantification Core (024614). This project has also been funded in part with federal funds from the National Cancer Institute, National Institutes of Health, currently under Contract No. 75N91024F00011. The content of this publication does not necessarily reflect the views or policies of the Department of Health and Human Services, nor does mention of trade names, commercial products, or organizations imply endorsement by the U.S. Government.

