## Supplemental metafile for "RhCMV Expands CCR5 Memory T Cells and promotes SIV reservoir genesis in the Gut Mucosa"

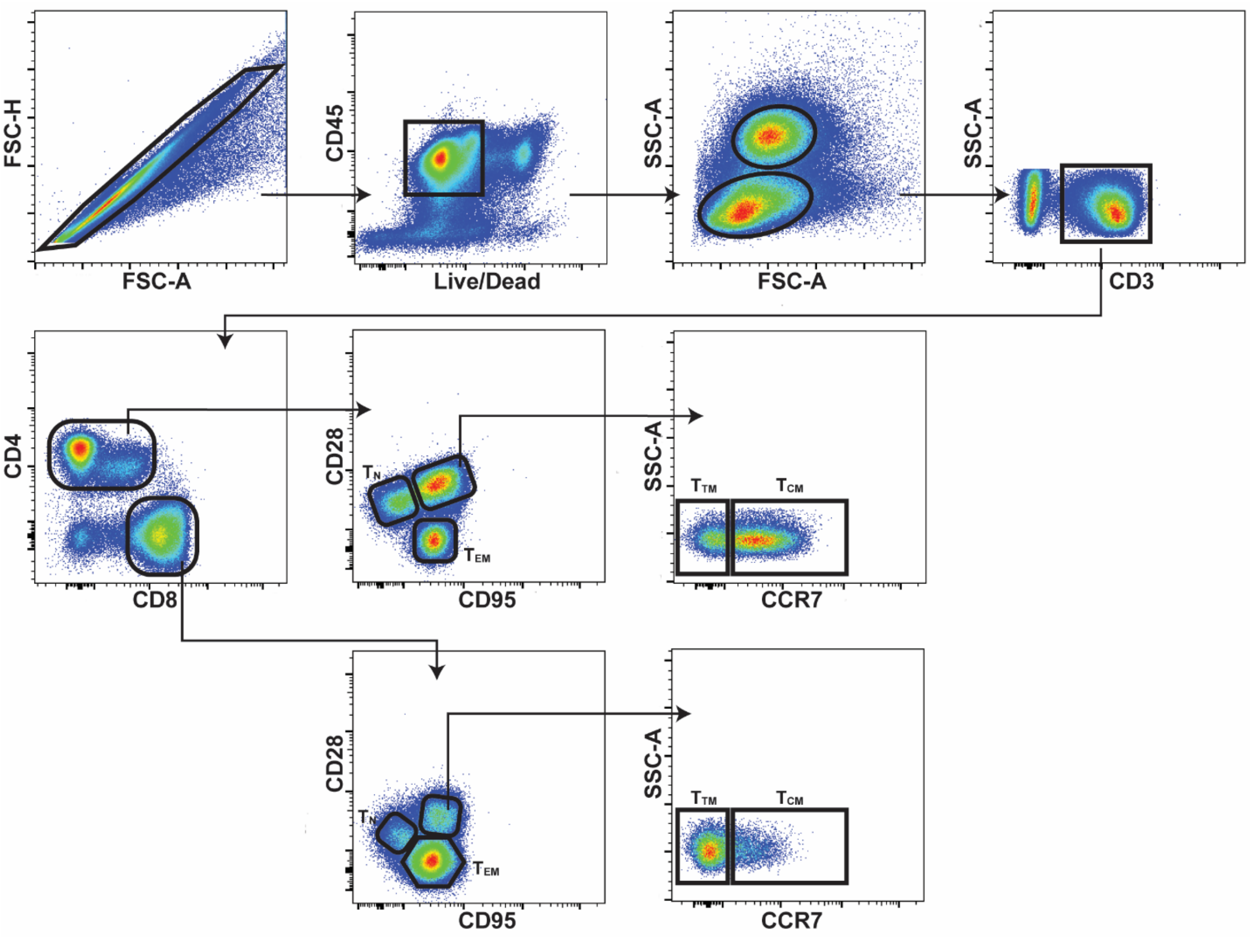


**Supplementary Figure 1:** Gating strategy for the CD4+ and CD8+ T cell maturation subpopulations. Key: $T_N$, naive; $T_{CM}$, central memory; $T_{TM}$, transitional memory; $T_{EM}$, effector memory.

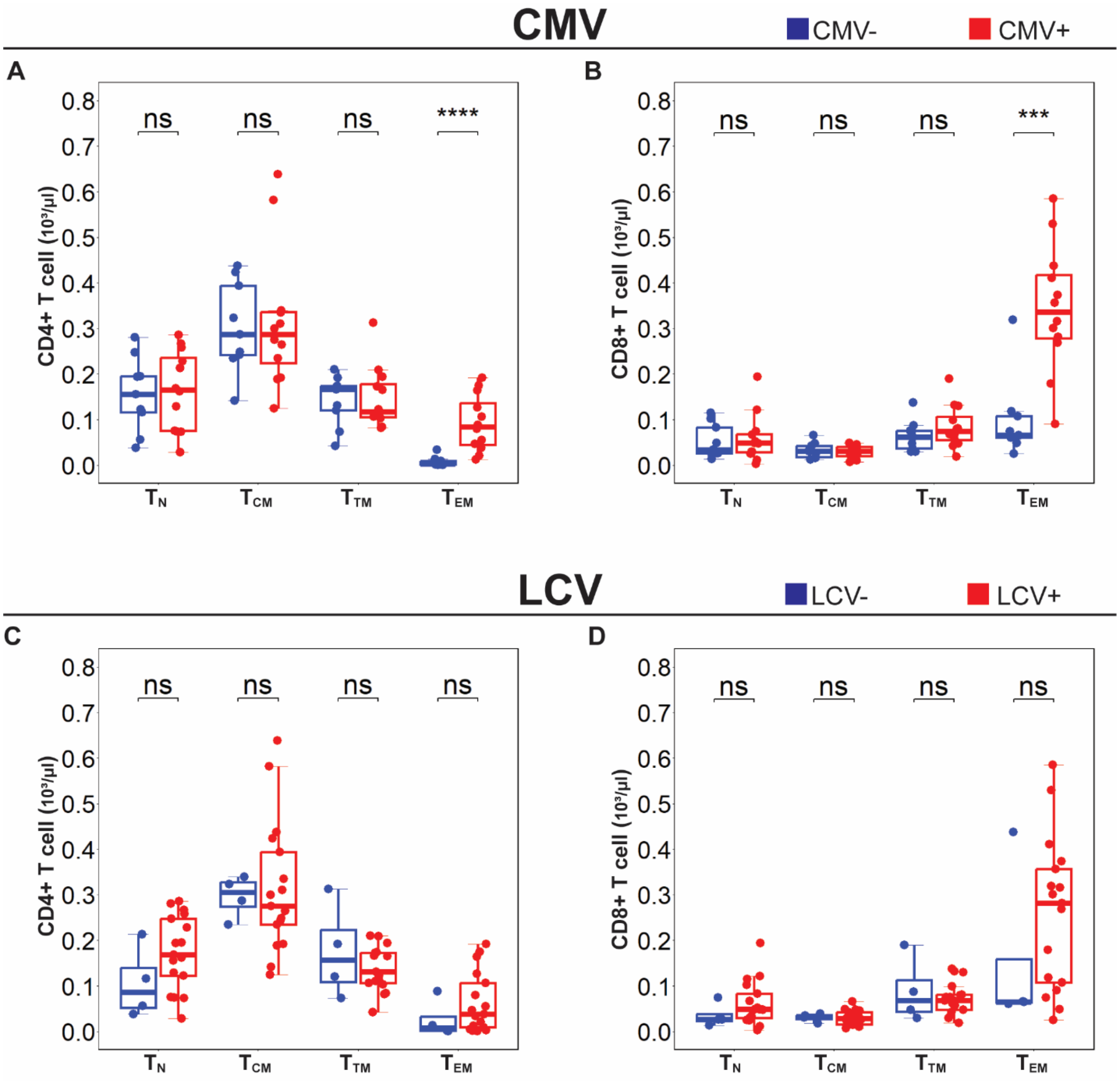


**Supplementary Figure 2:** Blood CD4+ (**A**) and CD8+ (**B**) T cell numbers across different maturation subpopulations by CMV-serostatus (CMV+ n = 12, CMV- n = 9). Blood CD4+ (**C**) and CD8+ (**D**) T cell numbers across different maturation subpopulations by LCV-serostatus (LCV+ n = 17, LCV- n = 4). Error bars represent 1.5 times the interquartile range. Statistical comparison performed using two-sided Mann-Whitney U test. Key: $T_N$, naive; $T_{CM}$, central memory; $T_{TM}$, transitional memory; $T_{EM}$, effector memory; ns $p>0.05$; *** $p<0.001$; **** $p<0.0001$.

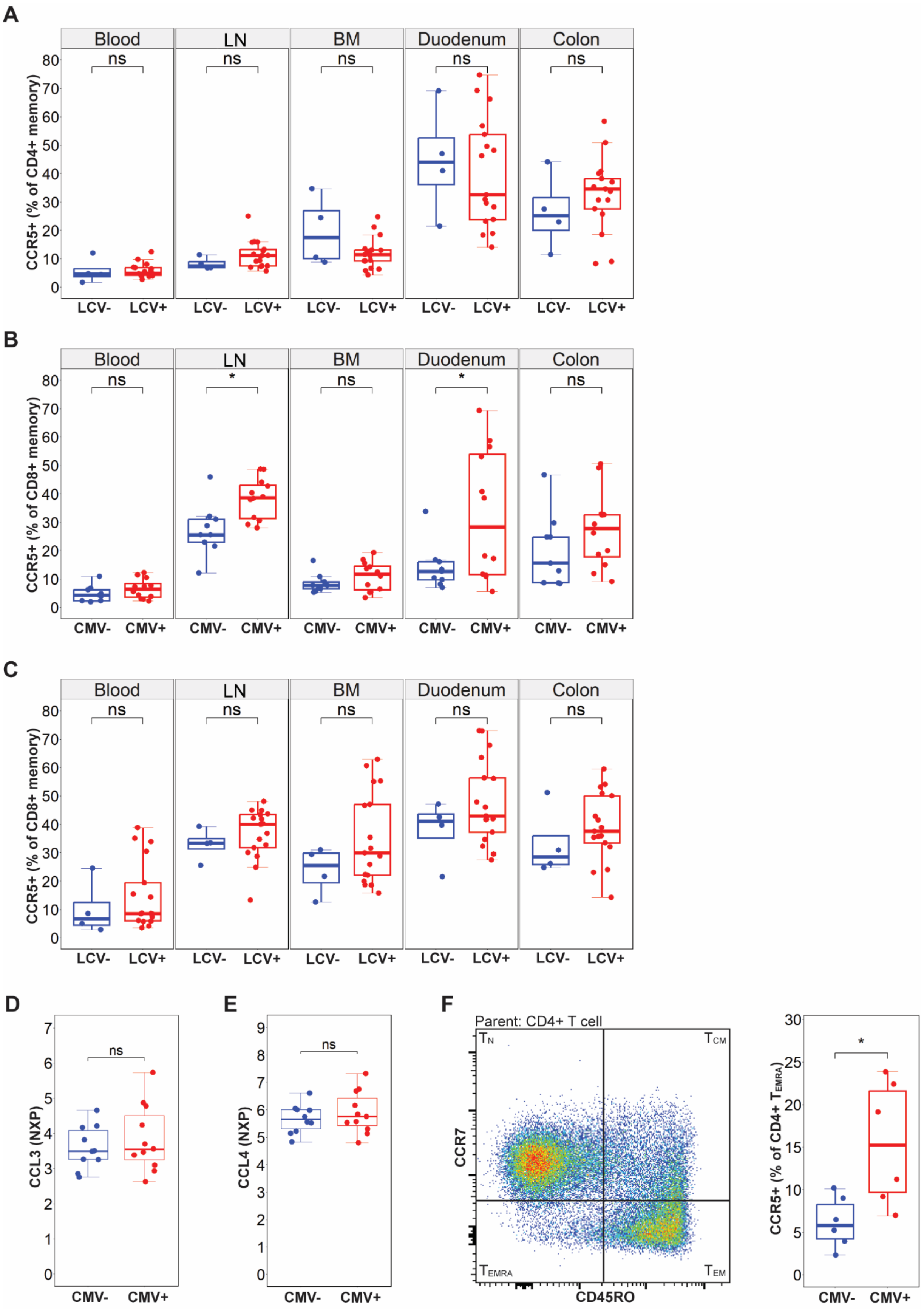
A
B
C
Blood
LN
BM
Duodenum
Colon
ns
*
CCR5+ (% of CD4+ memory)
CCR5+ (% of CD8+ memory)
LCV-
LCV+
CMV-
CMV+
D
E
F
CCL3 (NXP)
CCL4 (NXP)
Parent: CD4+ T cell
$T_N$
$T_{CM}$
$T_{EMRA}$
$T_{EM}$
CCR7
CD45RO
CCR5+ (% of CD4+ $T_{EMRA}$)

**Supplementary Figure 3: A** %CCR5+ CD4+ memory cells by LCV-serostatus. **B** %CCR5+ CD8+ memory cells by CMV-serostatus. **C** %CCR5+ CD8+ memory cells by LCV-serostatus. **D** Plasma CCL3 expression by CMV-serostatus. **E** Plasma CCL4 expression by CMV-serostatus. **F** Human PBMC CD4+ T cell maturation subpopulation gating strategy and %CCR5 CD4+ $T_{EMRA}$ cells. For all figures LCV+ n = 17, LCV- n = 4; CMV+ n = 12, CMV- n = 9. Error bars represent 1.5 times the interquartile range. Statistical comparison performed using two-sided Mann-Whitney U test. Key: LN, lymph node; BM, bone marrow; NPX, normalized protein expression; $T_N$, naive; $T_{CM}$, central memory; $T_{EM}$, effector memory $T_{EMRA}$, terminal effector memory; ns p>0.05; * p<0.05.

**A**

| Characteristics | All | CMV- | CMV+ | p-value |
|---|---|---|---|---|
| n | 110 | 35 | 75 | |
| Age (mean; years) | 12.31 | 12.26 | 12.34 | 0.79 |
| Age (min-max; years) | 1.93-18.01 | 7.39-17.41 | 1.93-18.01 | |
| Sex (M/F) | 67/43 | 21/14 | 46/29 | 1 |

**B**

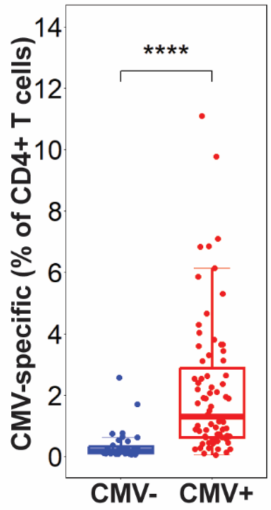


**Supplementary Figure 4: A** Age and sex characteristics of CMV+ and CMV- rhesus macaques from the large animal cohort. **B** %of CMV-specific CD4+ T cells by CMV-serostatus. Error bars represent 1.5 times the interquartile range. Statistical comparison performed using two-sided Mann-Whitney U test. Key: **** p<0.0001.

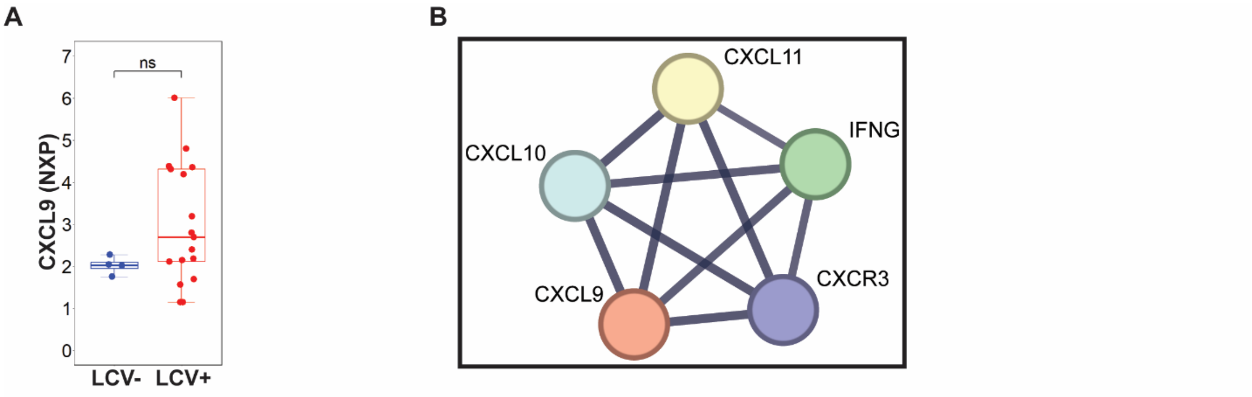


C

CD8+ Tem% Quartile Demographics

| Characteristics | All | Q1 | Q2 | Q3 | Q4 | p-value |
|---|---|---|---|---|---|---|
| n | 46 | 12 | 11 | 11 | 12 | |
| Age (mean; years) | 12.22 | 13.17 | 11.74 | 11.53 | 12.33 | 0.3013 |
| Age (min-max; years) | 8.28-17.43 | 8.40-15.63 | 8.28-17.43 | 9.28-15.28 | 8.55-15.38 | |
| CD8+ Tem% (mean) | 46.63 | 29.45 | 42.13 | 51.93 | 63.1 | <0.0001 |
| CD8+ Tem% (min-max) | 20.3-74.1 | 20.3-36.1 | 38.6-47.2 | 48.2-54.9 | 55.4-74.1 | |
| Sex(M/F) | 29/17 | 6/6 | 9/2 | 5/6 | 9/3 | 0.192 |

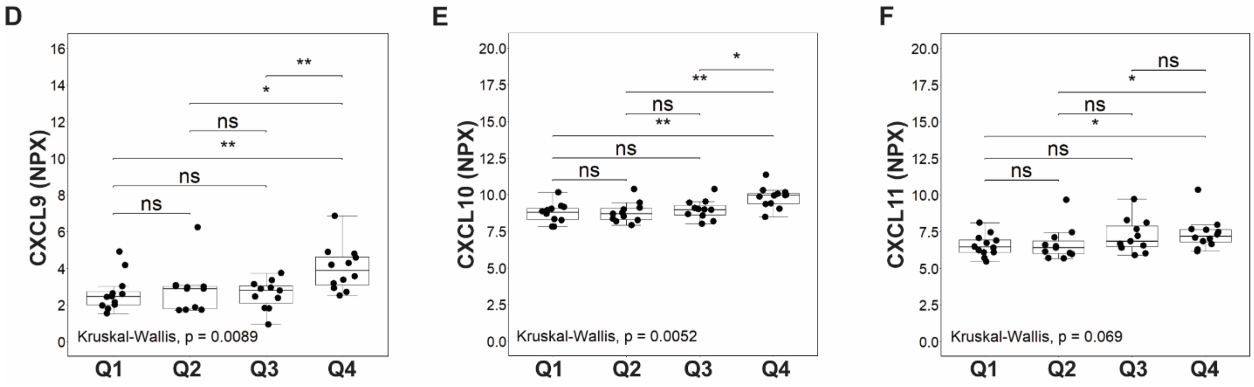


**Supplementary Figure 5: A** CXCL9 expression in the smaller cohort by LCV-serostatus. Statistical comparison performed using two-sided Mann-Whitney U test. **B** STRING network representing the interaction of IFNγ with CXCR3 and CXCL9/10/11. **C** Age and sex characteristics of CMV+ rhesus macaques from the large animal cohort divided by $\%T_{EM}$CD8+ T cells quartiles. **D-F** Expression of CXCL9/10/11 by $\%T_{EM}$ CD8+ T cells quartiles. Statistical comparison performed using two-sided Kruskal-Wallis test and post-hoc analysis by Dunn's test. Error bars represent 1.5 times the interquartile range. Key: ns p> 0.05; * p<0.05 ** p<0.001.

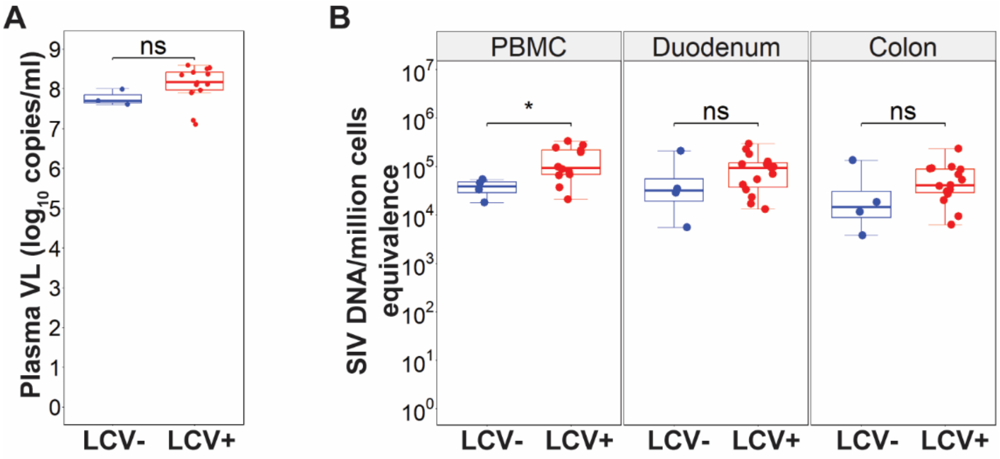


**Supplementary Figure 6: A** Plasma viral loads (VL) on day 13/14 by LCV-serostatus. **B** Cell-associated SIV DNA metrics by LCV-serostatus. For all graphs LCV+ n = 15; LCV- n = 4. Error bars represent 1.5 times the interquartile range. Statistical comparison performed using two-sided Mann-Whitney U test. Key: PBMC, peripheral blood mononuclear cells; ns p>0.05; * p<0.05.

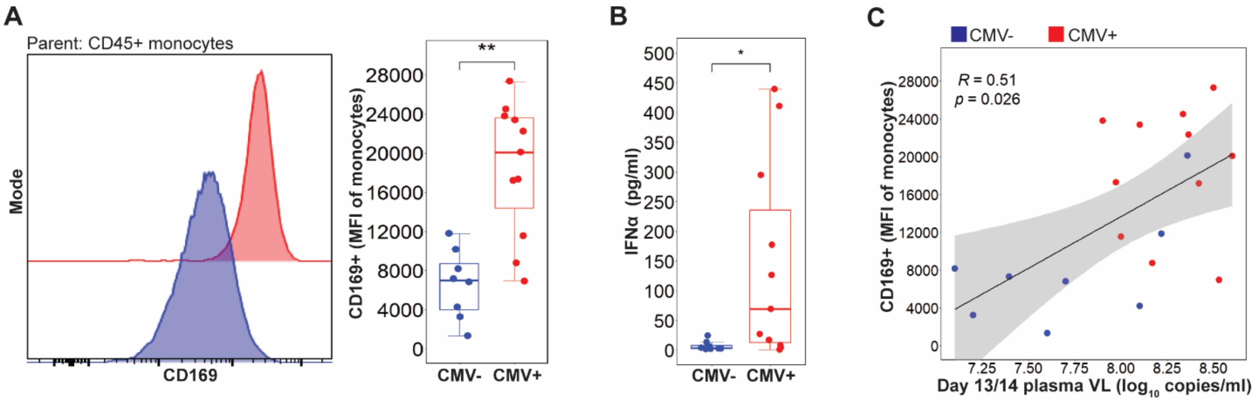


**Supplementary Figure 7: A** Histogram and CD169+ monocyte median fluorescent intensity (MFI) by CMV-serostatus in PBMC. **B** Plasma IFNα expression by CMV serostatus. **C** Two-sided Spearman's correlation between CD169+ monocyte MFI against plasma viral loads (VL). Shaded area represents 95% confidence interval. For all graphs CMV+ n = 11; CMV- n = 8. For A-B error bars represent 1.5 times the interquartile range. Statistical comparison performed using two-sided Mann-Whitney U test. Key: * p<0.05; ** p<0.01.

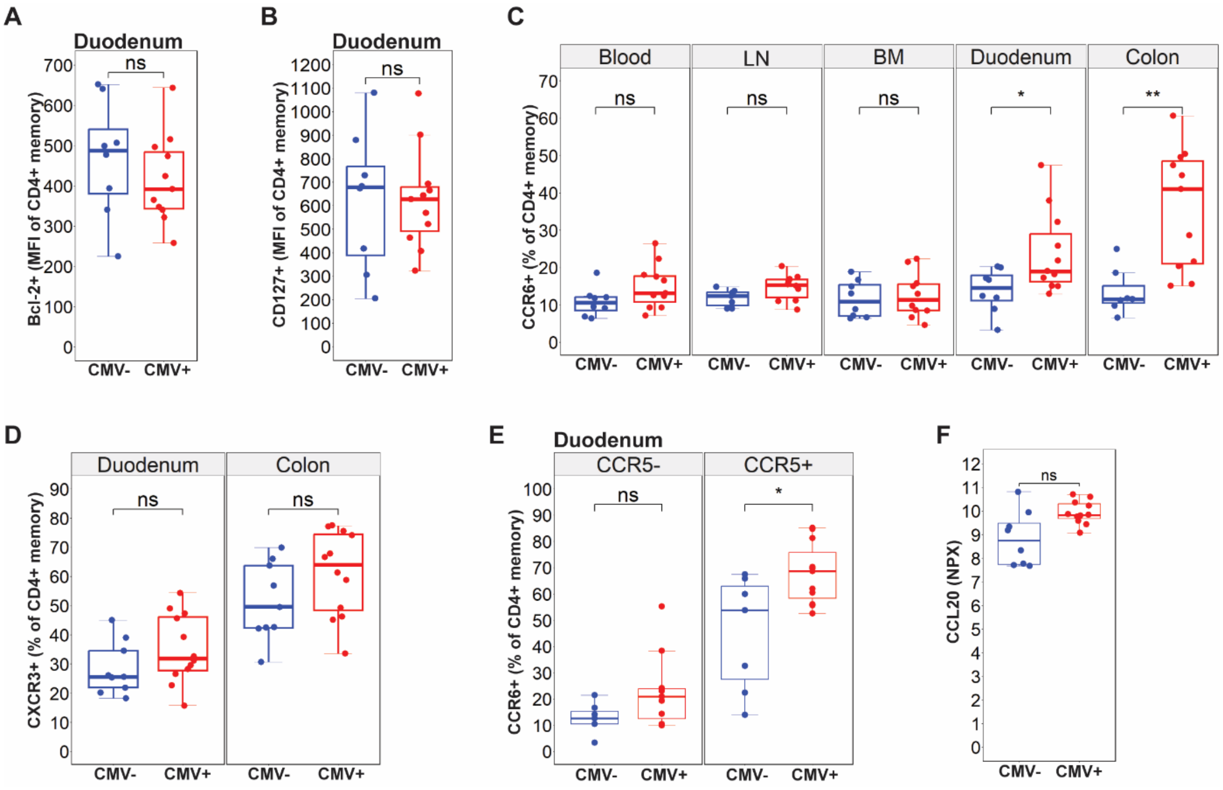


**Supplementary Figure 8: A** Bcl-2+ CD4+ memory median fluorescent intensity (MFI) by CMV-serostatus. **B** CD127+ CD4+ memory MFI by CMV-serostatus. **C** %CCR6+ CD4+ memory across sampled tissue by CMV-serostatus. **D** %CXCR3+ CD4+ memory across duodenum and colon by CMV-serostatus. **E** %CCR6+ CD4+ memory in CCR5+/- cells by CMV-serostatus. **F** Plasma CCL20 expression by CMV-serostatus. For all graphs CMV+ n = 11; CMV- n = 8. All data was taken during 13/14 dpi except D, at pre-infection. Error bars represent 1.5 times the interquartile range. Statistical comparison performed using two-sided Mann-Whitney U test. Key: LN, lymph node; BM, bone marrow; NPX, normalized protein expression; ns $p>0.05$; * $p<0.05$; ** $p<0.01$.
